# wgbstools: A computational suite for DNA methylation sequencing data representation, visualization, and analysis

**DOI:** 10.1101/2024.05.08.593132

**Authors:** Netanel Loyfer, Jonathan Rosenski, Tommy Kaplan

**Affiliations:** School of Computer Science and Engineering, The Hebrew University of Jerusalem, Israel; Faculty of Medicine, The Hebrew University of Jerusalem, Israel

**Keywords:** DNA methylation, Epigenetics, Computational biology, Liquid biopsy, Single-molecule fragment-level analysis

## Abstract

Next-generation methylation-aware sequencing of DNA sheds light on the fundamental role of methylation in cellular function in health and disease. These data are commonly represented at a single CpG resolution, while single-molecule fragment-level analysis is often overlooked.

Here, we present *wgbstools*, an extensive computational suite tailored for methylation sequencing data. wgbstools allows fast access and ultra-compact anonymized representation of high-throughput methylome data, obtained through various library preparation and sequencing methods. Additionally, *wgbstools* contains state-of-the-art algorithms for genomic segmentation, biomarker identification, genetic and epigenetic data integration, and more. wgbstools offers fragment-level analysis and informative visualizations, across multiple genomic regions and samples.

## Background

DNA methylation is an essential component of gene regulation and genome packaging and is a fundamental mark of cell identity. Indeed, alterations and dysregulation of DNA methylation are implicated in development, aging, and multiple diseases [1]. Analysis of DNA methylation is therefore key for understanding biological and developmental processes, as well as for clinical diagnosis. Notably, methylation-based analysis of circulating cell-free DNA fragments shed into the bloodstream following remote cell death events, is emerging as an accurate tool for quantifying cell-type-specific damage in clinical diagnosis and monitoring [2–8].

Genomic studies of DNA methylation often use the Illumina BeadChip 450K/EPIC arrays, which measure the average DNA methylation levels (beta values) at a predefined set of CpG sites, including a mere 1.5-3% of the 28 million CpG methylation sites across the entire human genome [9,10]. Recent advances in DNA sequencing technologies have facilitated a fragment-centered view that captures binary DNA methylation patterns of multiple neighboring CpG sites, at a single-molecule resolution [4–8,11–14]. These technologies include conversion methods such as bisulfite [15] or enzymatic methyl treatment followed by sequencing (EM-seq) [16], alongside direct detection of base modifications, using methylation-aware DNA sequencing technologies such as Oxford Nanopore Technologies (ONT) [17] or PacBio [18]. DNA methylation information could be measured across the entire genome, or enriched at target regions using hybrid capture arrays, restriction enzymes (RRBS), or targeted PCR [3– 5,19–21]. Nonetheless, the computational and algorithmic tools for processing, visualizing, and analyzing such data lag behind.

Here, we present wgbstools (github.com/nloyfer/wgbs_tools), an open-source computational suite for efficient end-to-end processing, data conversion, representation, anonymization, visualization, and analysis of sequencing DNA methylation data, across multiple sequencing platforms and library preparation methods.

## Results

### Data Representation

The primary format utilized for DNA methylation sequencing data is the BAM file format, which encompasses sequenced reads, names, genomic mapping, quality, and read pair information [22]. However, working with these files can be challenging due to their considerable size and complexity, making simultaneous analysis of multiple samples arduous. Furthermore, BAM files store complete genetic information and are bound by stringent privacy regulations that restrict data sharing and accessibility [23].

Consequently, bed/bigwig formats have become the prevailing method of representing DNA methylation data. These formats store the average methylation levels of individual CpG sites or the counts of methylated and unmethylated CpGs. Although these formats are more compact, they sacrifice fragment-level information, including the binary methylation patterns of neighboring CpGs within the same DNA fragment. The interdependencies among CpGs within a single fragment encompass crucial aspects of DNA methylation-based analysis, particularly for tissue-of-origin inference, in both health and disease [4–7,24].

wgbstools offers a solution for converting DNA methylation data from BAM format into the compact and indexed PAT format (Figure 1). This conversion involves merging read pairs, discarding all non-CpG positions, sorting by position, collating independent fragments that cover the same CpG sites and show identical methylation patterns, and indexing the data at the fragment level, resulting in a methylation-lossless compressed format, 300-fold more compact in disk size (Supplemental Table S1). The PAT format is founded on principles similar to those of the epi-read format introduced by *DNMTools* [25] and *biscuit [26]*, yet it is distinguished by a complementary set of utilities. These utilities encompass visualization, clipping, aggregation, filtering by the number of covered CpGs or using epigenetic fragment-level data and/or genetic information, and other commonly employed features (Supplemental Table S2).

**Figure 1.**
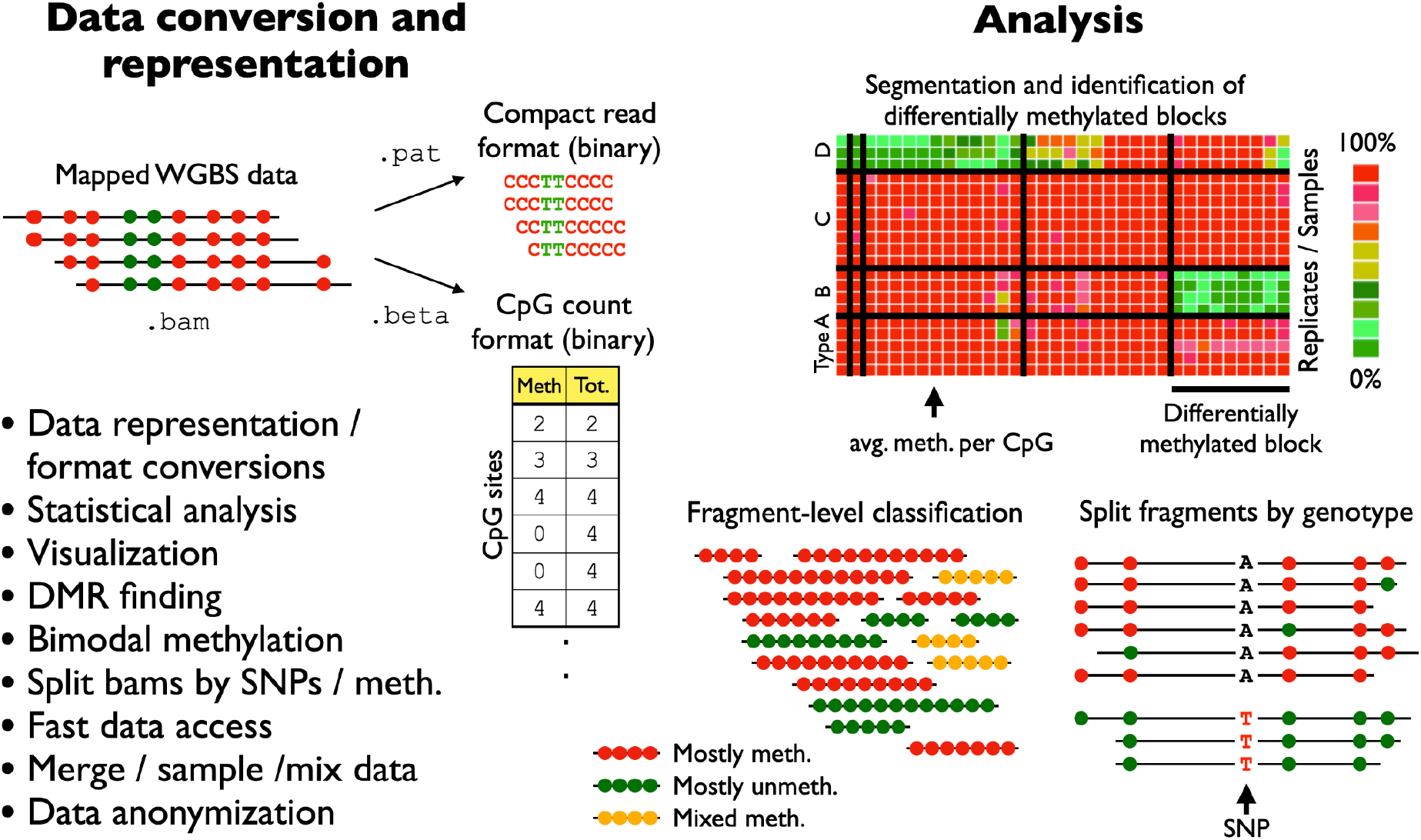
wgbstools: a software suite for DNA methylation data representation, analysis and visualization. Given a BAM file containing next-generation sequencing of DNA methylation data, wgbstools converts mapped reads (or read-pairs) to compact representations of methylation data at fragment-level resolution, thus maintaining methylation patterns across adjacent CpG sites from a single molecule. Additionally, wgbstools provides a CpG-oriented count format which includes the number of reads in which any CpG was methylated and sequenced. All wgbstools formats are fully indexed to support efficient direct access and include multiple visualization schemes, for command line explorations as well as in publication quality. wgbstools contains several machine learning algorithms for optimal segmentation of genome-scale data into continuous methylation blocks of dynamic size; for the identification of cell-type-specific differentially methylated regions; and SNP-based allele-specific methylation data analysis.

wgbstools also supports the CpG-centered binary format BETA, which stores the total number of reads covering each CpG, and how often it was methylated (Figure 1). Files of this format use a fixed compact size of ∼50Mb, allowing efficient direct access for target CpGs, simultaneously done across hundreds of samples.

### Analysis and Visualization

wgbstools provides extensive support for data manipulation and visualization, both for command-line explorations and for figure generation. Users can efficiently perform tasks such as read sub-sampling for simulating different sequencing depths, merging replicates, or simulating in silico admixtures by combining files at various concentrations.

We have also developed a statistical framework that enables the identification of genomic regions exhibiting bimodal methylation patterns, by which half of the sequenced fragments are typically methylated, whereas the other half are mostly unmethylated, for example in regions of allele-specific methylation due to parental imprinting or meQTL genetic variation. These regions can be further explored using wgbstools’ `split_by_meth` and `split_by_allele` features, which select specific reads from an input BAM file, based on their fragment-level methylation/genotype at a target single nucleotide polymorphism (SNP), thus facilitating analysis of allele-specific methylation and parental imprinting (listed below).

wgbstools offers a range of informative visualizations at both fragment-level and single-CpG resolutions, enabling the examination of binary patterns and average methylation across multiple files. Overall, wgbstools presents a comprehensive command-line suite that encompasses data processing, analysis, anonymization, visualization, and manipulation tasks. It can be used independently or in conjunction with other tools such as the UCSC genome browser [27], IGV [28], or biscuit [26] for enhanced functionality.

### .pat and .beta - fragment-level and CpG-oriented data formats

BAM file information is analyzed and converted into two data formats, at complementary levels of representation and compactness. The first format, called PAT, preserves the fragment-level DNA methylation data via the `bam2pat` command. First, the methylation sites in the reference genome are indexed as CpG_1_, CpG_2_, CpG_3_, …, CpG_N_ with N=28,217,448 for hg19. Each sequenced read is then projected onto this N-dimensional space, by discarding non-CpG positions. To further conserve space, sequenced reads (or read-pairs) that cover the same set of CpG sites and show an overall identical methylation pattern are merged and counted together (Figure 2). PAT files are compressed using *bgzip* and indexed for rapid direct access using *tabix* [29].

**Figure 2.**
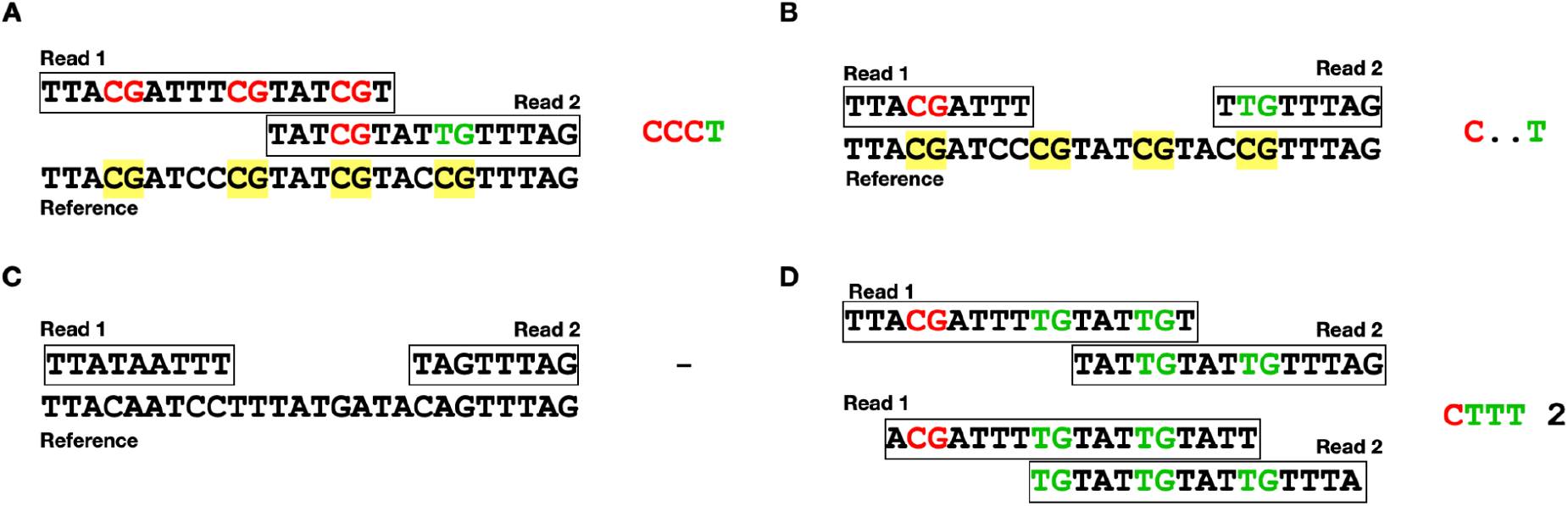
Interpretation of methylated DNA sequences. `bam2pat` infers methylation state for each sequenced CpG site by comparing sequenced reads to reference genome, and merges overlapping reads to produce the binary methylation pattern (PAT format). (A) Overlapping read pair, aligned against the reference genome (CpG sites marked in yellow). Following bisulfite or enzymatic conversion, all cytosines are converted and read as T, whereas methylated cytosines are protected and read as C. Shown are three methylated CpGs, followed by an unmethylated one (denoted by “CCCT” in the PAT file). (B) Same as (A), for a non-overlapping read-pair. The methylation status of the two center CpGs is unknown, and denoted as a “C..T” pattern. (C) A sequenced read-pair that overlaps no CpG sites, and is discarded from the PAT file. (D) DNA fragments, showing the same binary pattern across the same CpG sites are merged (and counted) for compactness (e.g. “CTTT 2”)

The second format, called BETA, retains methylation counts for individual CpGs. These files are of fixed size, equivalent to the size of the reference genome. For each CpG, two numbers are stored as unsigned integers (8/16 bits): the count of methylated observations, and the total count of observations (Figure 1). This format only keeps CpG information (as opposed to fragment-level information which is kept in PAT files), to prioritize compression efficiency and runtime. The beta files enable ultra-fast reading, writing, and direct access, making them particularly advantageous for aggregating and integrating information from a large number of samples.

Below, we describe the main features of wgbstools:

### The view command

The `view` command allows users to retrieve and display all fragments from a PAT file, within a specified genomic region, a set of regions, or across the entire genome. The command also provides various filtering options, including fragment length (i.e. number of covered CpGs), fragment clipping to specific boundaries, fragment sub-sampling, and more. When applied to a BETA file, the `view` command displays the per-CpG binary methylation information, as well as chromosomal coordinates, in BED format [27]. Overall, the `view` command provides a flexible and customizable approach for viewing and filtering methylation data, enabling users to extract specific subsets of data for further analysis and exploration.

### The vis and pat_fig commands

The `vis` command offers in-terminal visualizations for one or more samples in parallel, focusing on a target genomic region. This command utilizes ANSI colors and symbols for clear lightweight visually compelling graphics, printed directly to the terminal for interactive explorations (Figure 3).

**Figure 3.**
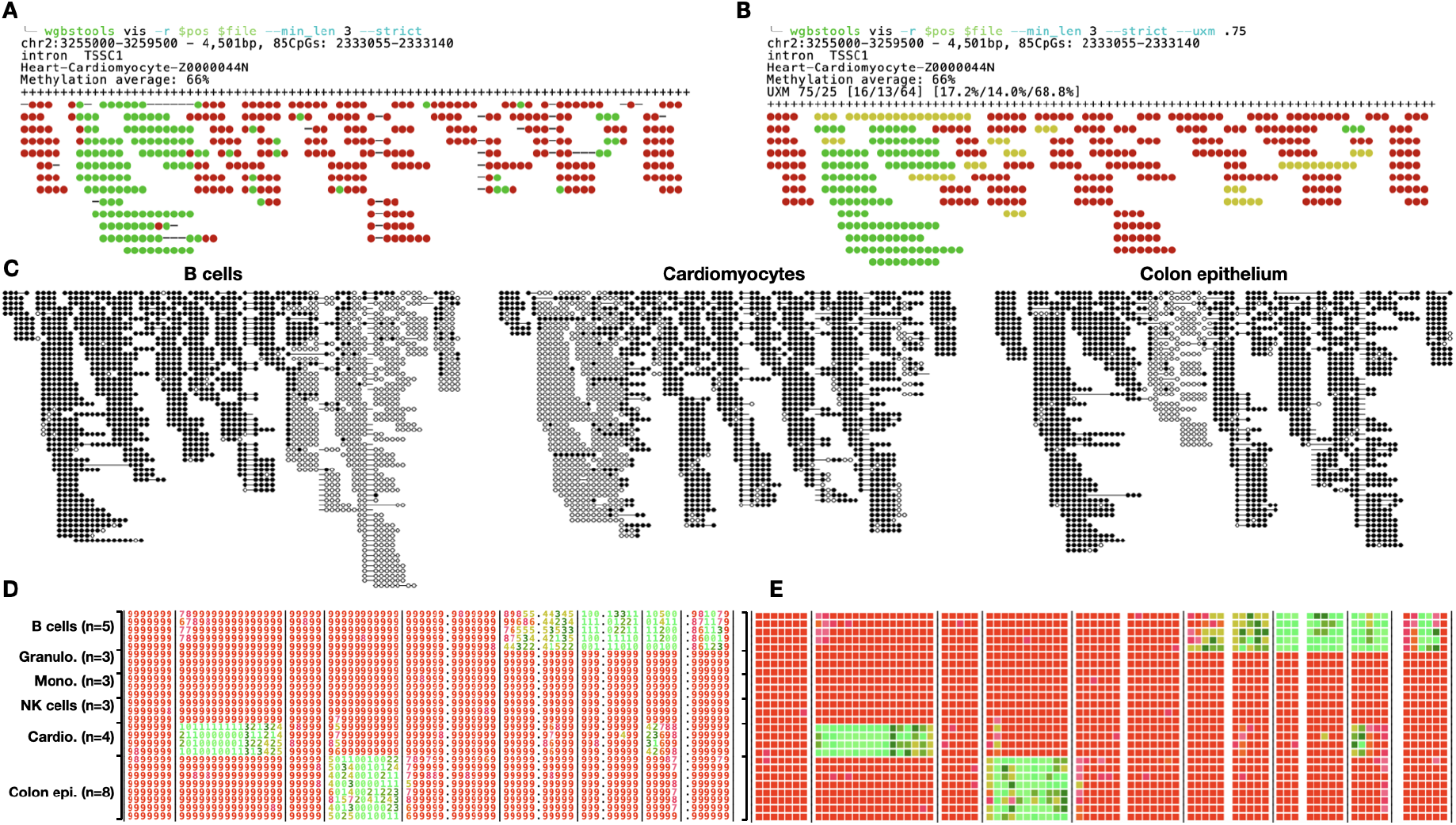
wgbstools’ vis command. **(A)** A fragment-level command-line interface for the visualization of a PAT file. Each individual string shows one sequenced fragment, whereas red circles denote methylated CpGs and green unmethylated ones. Strikethrough positions denote missing CpGs (e.g. in non-overlapping paired-end reads from longer DNA fragments). Fragments are stacked and aligned by mapped positions. Shown are 85 CpGs from a 4.5Kb region from purified cardiomyocytes. **(B)** Same as (A), where fragments are classified by their overall fragment-level methylation state (green: ≤25% methylated CpGs; red: ≥75%; otherwise yellow). **(C)** PDF visualization of multiple BETA files, for the same genomic region, including cardiomyocytes, colon epithelial cells, B cells, all from Loyfer et al. **(D)** Screenshot of command-line interface, showing average methylation per CpG/sample in 0-9 range, in color. **(E)** same as (D), using the --heatmap flag, and the --blocks option, which uses vertical bars based on genomic segmentation of homogeneously methylated regions.

When applied to PAT files, the `vis` command displays a stacked view of aligned fragments from this target region. Each fragment is visualized as a continuous string of CpGs, colored based on their methylation. Optional features include the ability to filter reads by length, clip reads to the specific region, classify and color reads based on their overall average methylation, randomly shuffle read order, print genomic boundaries, and print regional statistics and annotations.

Similarly, `pat_fig` generates high-quality publication-ready plots of methylation data from PAT files, in a variety of file formats (e.g. pdf, png), as well as multiple esthetic customizations (plot size, color schemes, spacing, etc.; see Figure 3).

When applied to BETA files, the `vis` command computes the average methylation for each CpG in each sample and prints a colorful heatmap. For compactness, average methylation values are discretized into values ranging from 0 to 9, represented by a single colored ASCII character. This approach allows for an informative and visually appealing visualization of the methylation patterns in the selected CpG sites.

As shown in Figure 3, the `vis` command provides a convenient and visually intuitive way to explore and analyze methylation data directly in the terminal, enabling users to gain valuable insights and make informed interpretations of their data.

### The segment and beta_to_table commands

Neighboring CpGs often display correlated methylation patterns in a block-like fashion [4–6]. The `segment` uses a computational, dynamic programming algorithm, to find the optimal segmentation of the genome into contiguous regions of neighboring CpGs that show similar methylation levels across samples (Fig. 3E) [5]. The output of this command is a BED file, where the genomic coordinates and CpG IDs of each block are denoted. Parameters include a Bayesian pseudocount value, which controls the tradeoff between block length and homogeneity; a list of BETA files on which the (concurrent multi-channel) segmentation is performed; and thresholds for minimal and maximal block sizes (in bases or CpGs).

Once a block file is obtained, the `beta_to_table` command processes a set of samples (in BETA format) and produces a tab-separated table showing the average methylation of each region (row) across each sample (column). Given an additional “groups” file, the command allows aggregation over replicates or samples of the same cell type. To reduce noise, the program can treat low-coverage regions, where few reads are present, as missing values.

`beta_to_table` is applicable to any BED file, including custom cell-type-specific markers, differentially methylated regions, or other regions of interest in the genome. This segmentation and block representation allows for an easy dimensionality reduction, moving from the 28 million CpG space to a compact biologically meaningful representation of whole-genome DNA methylation.

### The find_markers command

The `find_markers` command identifies differentially methylated blocks (DMBs) that exhibit unique methylation patterns in one or more samples from a specific cell type or condition (vs. all or some others). For instance, in a dataset consisting of k distinct cell types, each in multiple replicates, the command identifies and ranks the top genomic regions (blocks) that are specifically unmethylated in most replicates of one cell type, compared to the samples in the remaining k-1 cell types (one-vs-all approach).

`find_markers` requires users to input an initial set of blocks (e.g. a capture-panel design file, a set of promoters or other regions of interest, or regions found using the `segment` command), as well as a collection of samples to be compared (BETA format), and a “groups” file assigning each sample to its respective group / cell type. Markers are scored and ranked using several possible scores, and filtered by statistical significance (Figure 4, Methods) [5].

**Figure 4.**
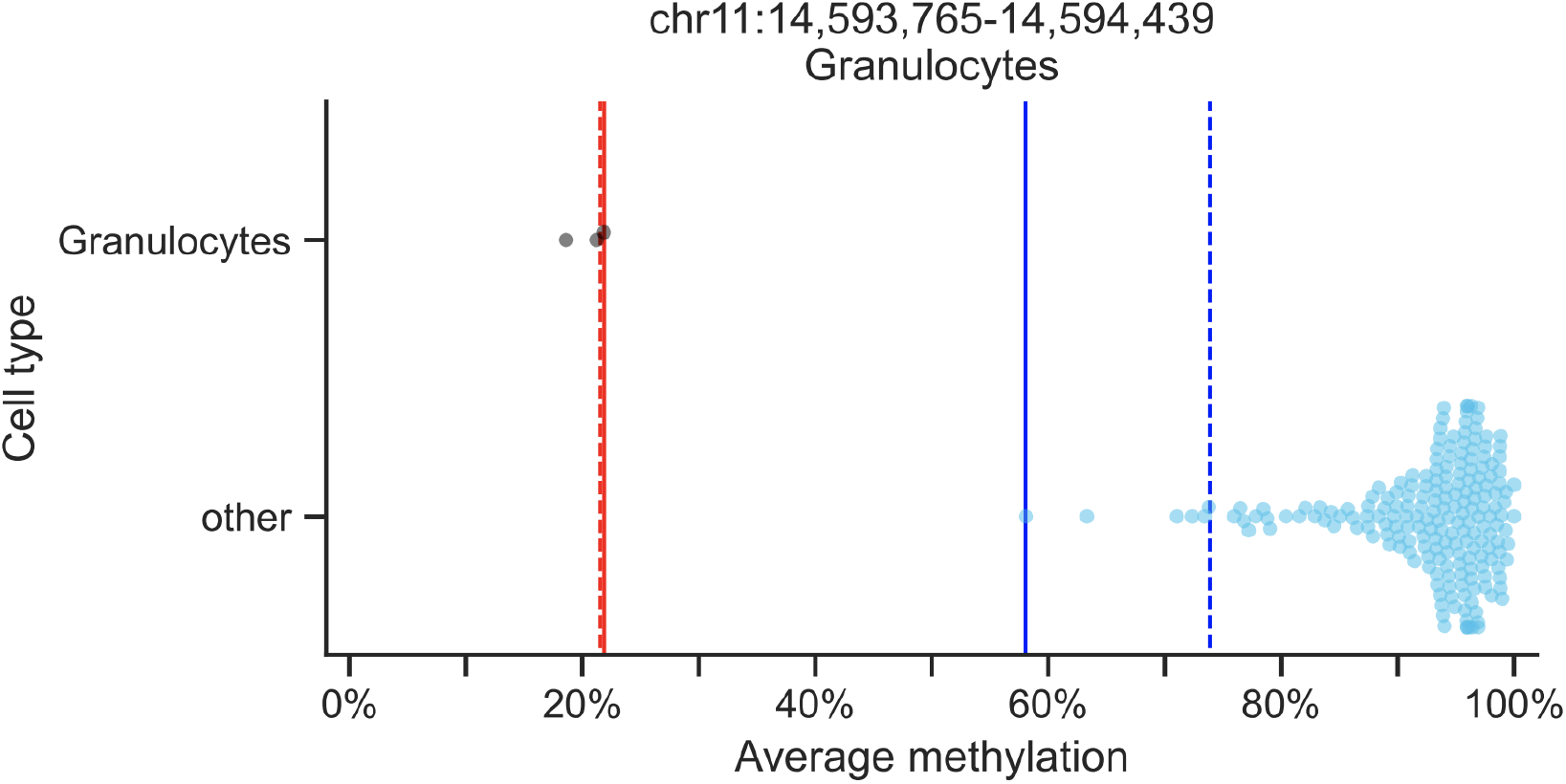
Marker finding specification. Shown the average methylation values of 205 WGBS samples from Loyfer et al. in a block of 5 CpGs (chr11:14,593,765-14,594,439, 674bp, hg19). `find_markers` may identify this block as a marker for granulocyte cells as the “target group” since the methylation values are low in the granulocyte samples (black circles, n=3) and high in the other samples (light blue circles, n=202). `find_markers` tests the values of, and the difference between, the 2.5^th^ percentile of the background group (dashed blue line) and the 75^th^ percentile of the target group (dashed red), as well as the difference between the minimum of the background group (solid blue) and the maximum of the target group (solid red). The T-test p-value (2.8e-46) is reported and is also used by `find_markers` for block filtering.

**Figure 5.**
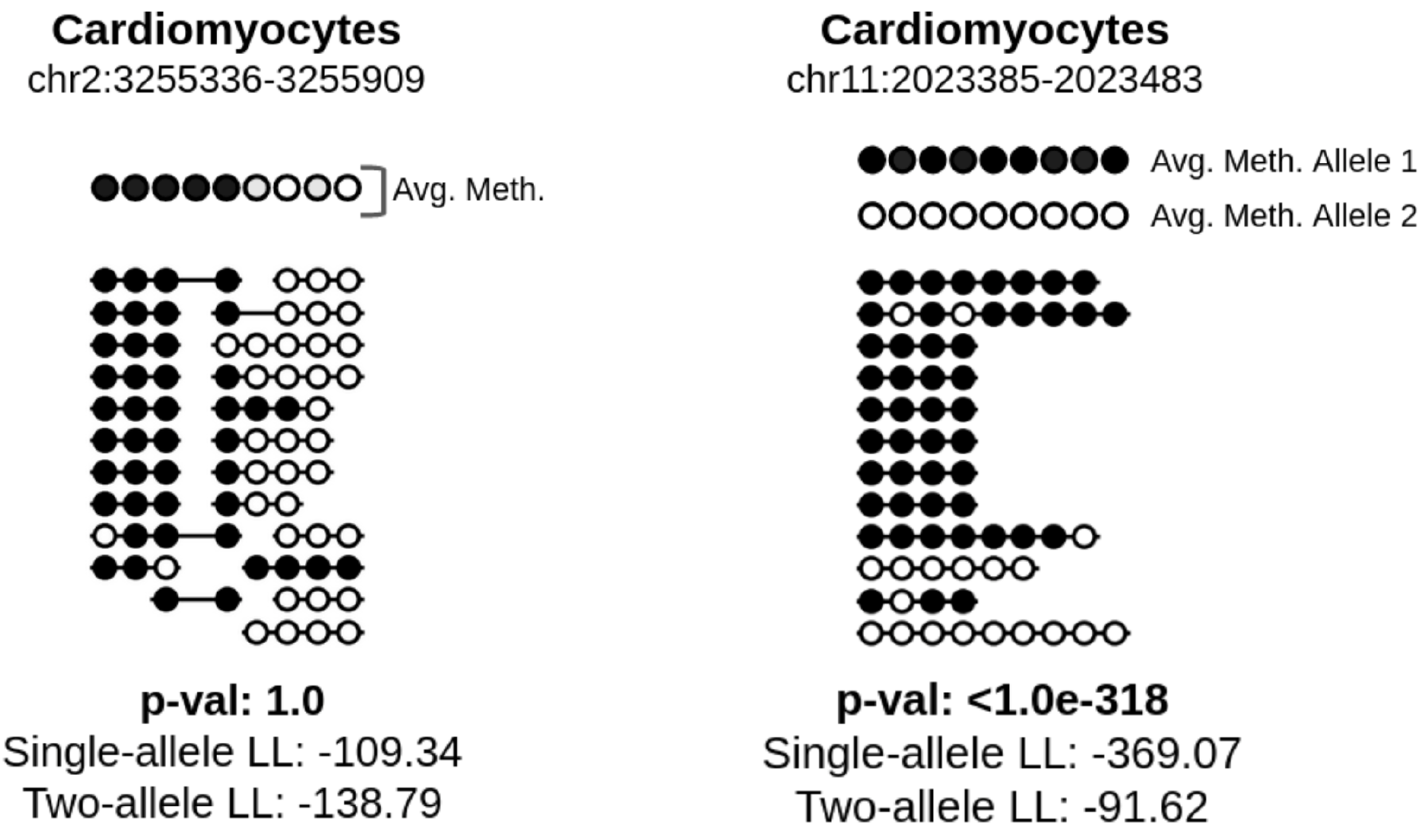
test_bimodal function identifies bimodal regions with significant p-values. We show methylation of CpG sites on sequenced fragments from a single cardiomyocyte sample at two distinct regions. The first region, on the left, shows a region that is mostly methylated and then becomes mostly unmethylated, however each CpG site has the same methylation pattern across all sequenced fragments, and is thus not bimodal and receives an insignificant p-value. The second region, on the right, is from a known imprinting control region, and each CpG site is either methylated or unmethylated depending on whether it originated from the paternal or maternal allele. For this reason, a probabilistic model which allows for two distinct alleles each with differing probabilities for each CpG site to be methylated is more likely than a single-allele model.

### The pat2beta command

The `pat2beta` command is designed to process a pat file and generate a corresponding beta file. The resulting beta file stores, for each CpG site, the number of reads covering the site and the number of observed methylated instances of the site (Figure 1). This command creates a compact representation of marginal methylation data in the beta file format.

### The convert command

The `convert` command provides functionality for translating genomic loci between CpG indexes and genomic coordinates. In wgbstools, genomic regions are indexed using CpG-based coordinates, denoted as CpG_1_, CpG_2_, CpG_3_, and so on. This indexing approach optimizes memory usage and running time. The `convert` command takes a bed file as input and adds columns that indicate the CpGs covered by each line in the bed file. Moreover, it appends genomic annotations for each region, providing valuable context and information for the converted data.

### The homog command

This command is designed for fragment-level analysis of next-generation sequencing DNA methylation data. Given a PAT file, and an input BED file containing a set of regions of interest, `homog` classifies each sequenced fragment as mostly unmethylated (U), mostly methylated (M), or mixed (X), and outputs for each genomic regions the number U, X, and M fragments. The BED file must include at least 5 columns: chromosome, start, end, start-CpG-index, and end-CpG-index, as outputted by the `convert` command. By default, `homog` considers fragments covering ≥3 CpGs, and defines U fragments as those having ≤ 25% methylated CpG sites, and M fragments as those with ≥ 75% methylated CpGs. For efficiency and compactness, the default output file is in a binary uint8 format, although tab-delimited text output is also available.

### The test_bimodal command

This command implements a statistical test to estimate whether the sequenced fragments in a given target region are likely to have been derived from a single methylation pattern (i.e. biallelic pattern), or from a two-component mixture model, representing differential, allele-specific methylation. The null hypothesis H_0_ assumes each of the CpGs in the target region is independent of its neighboring sites, and a specific Bernoulli distribution parameter is fitted using a maximum likelihood estimator for each CpG site. Conversely, the H_1_ hypothesis allows two such distributions, each occurring with a prior probability of 50% (Methods, Supplemental. Figure S1).

## Discussion

The compact nature of the formats generated by wgbstools significantly reduces the disk space and memory requirements for analyzing DNA methylation sequencing data. This is particularly crucial in the face of the escalating volume of data generated by next-generation sequencing technologies and the increasing popularity of DNA methylation studies, allowing for joint analysis and interpretation of multiple samples. The ease of downstream analysis is thereby enhanced, allowing researchers to handle larger datasets more efficiently, and to foster the exploration of complex relationships within methylation patterns. wgbstools’ PAT format is used by the UXM deconvolution method [5], a computational fragment-level reference-based computational algorithm for DNA methylation sequencing data, that is useful for identifying of cell-of-origin in liquid biopsy, in computational pathology, etc.

wgbstools introduces a paradigm shift in the sharing of methylation sequencing data. Today, researchers commonly release derivative formats (beds/bigWigs) of their data publicly, while withholding or restricting access to the raw sequencing data due to its sensitive nature containing private information about donors. In contrast, our presented PAT format achieves a delicate balance, revealing only the methylation pattern, while discarding the DNA sequence, which holds the more sensitive information. This anonymization proves to be a significant asset, particularly for large-scale human methylation projects like Roadmap Epigenomics [30], BLUEPRINT [31] and the comprehensive human WGBS atlas by Loyfer et al. [5]. By providing an ultra-compact CpG-oriented representation of high-throughput data, researchers can now share methylation data without compromising privacy, facilitating broader collaboration and accelerating scientific discoveries.

While wgbstools excels in CpG-specific resolution, it does come with a trade-off. The tool prioritizes CpG sites, potentially neglecting valuable information such as motifs or fragment start/ends crucial for fragmentomic analysis. Researchers interested in retaining such information while leveraging some of wgbstools features could utilize wgbstools’ `add_cpg_counts` feature, which adds the methylation pattern (e.g. CCCTCTCTT) as an additional tag for each read-pair in the BAM file, allowing users to avoid fully transitioning to the PAT format domain.

wgbstools offers universal compatibility by supporting any BAM file, irrespective of the sequencing technology employed, be it Illumina, Ultima Genomics, Oxford Nanopore, or others. This flexibility ensures that researchers can seamlessly integrate data from diverse sources, promoting cross-platform comparisons and enhancing the robustness of methylation analyses.

## Conclusions

As sequencing-based DNA methylation data analysis is gaining popularity and is rapidly evolving, there is a growing need for software for data processing, representation, visualization, and analysis. wgbstools is a flexible suite that allows researchers from the biomedical community to analyze and visualize DNA methylation sequencing data efficiently and at ease, and will pave the way for novel discoveries, including cell type-specific enhancers, novel imprinting regions, and novel methylation-based biomarkers for circulating cell-free DNA.

## Methods

### Differentially methylated block identification method

The `find_markers` command essentially filters a set of blocks to a subset of differentially methylated ones, by selecting blocks where the average methylation value of the samples in one set (“target” group) is different from the values in the other set of samples (“background” group). Multiple statistics are computed for each block and each group, such as the average of averages, the k^th^ percentile, minimum/maximum, and the percentage of missing values in each group. It then filters the blocks by multiple conditions, including differences in the statistics between the two groups, absolute thresholds of the statistics, significance level of a T-test, direction (i.e., methylated vs. unmethylated), block size, and number of CpGs. This allows the user to filter and sort the markers by these statistics, the differences between them, and a t-test p-value. See Figure 4 for an example.

### Test_bimodal

We calculate a p-value for the significance of a two-allele model being more likely than a single-allele model for a specific region. We assume that each CpG site has its own probability of being methylated. That is, for CpG site *i* the probability to be methylated is θ_*i*_, and any read intersecting this site has probability θ_*i*_ to be methylated at that CpG site. Given the probability of being methylated, we assume the methylation status of each CpG site in each read is independent of the status at other sites. If we let *r*^*j*^ be read *j* and 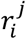 be the methylation status of read *j* at CpG site *i*, where it equals 1 if it is methylated and 0 otherwise. For the single-allele model, the likelihood can then be written as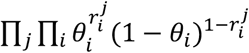. We use the maximum likelihood estimator for the probability to be methylated at each CpG site (# methylated observations / total observations).

For the two-allele model, We let A be a variable representing whether a read is distributed according to allele 1 or allele 2. Thus the likelihood can be written as:

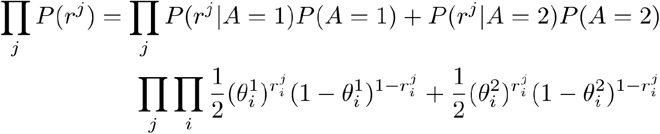

This can be seen in supplementary figure 2. We use an expectation-maximization (EM) algorithm to calculate the likelihood by iteratively associating reads with an allele and estimate the probability to be methylated at each CpG site for each allele. Finally, we calculate a p-value using the log-likelihood ratio test to ensure that the two-allele model significantly describes the data better than the one-allele model.

### System requirements

wgbstools functions exclusively on Unix-based systems and is not compatible with Windows. Leveraging standard C++ libraries, widely-used Python packages, and established bioinformatics software such as SAMtools[19505943], wgbstools is easy to install. Specific features may necessitate additional Python packages; users can find detailed requirements on the project’s GitHub page.

## Abbreviations

WGBS: whole-genome bisulfite sequencing
RRBS: reduced representation bisulfite-sequencing
EM-seq: enzymatic methylation sequencing
DMR: differentially methylated regions
DMB: differentially methylated blocks
PCR: polymerase chain Reaction

## Availability of data and materials

The source code is available at https://github.com/nloyfer/wgbs_tools. This code is distributed under the Hebrew University of Jerusalem open software research license. The license agreement is available at https://github.com/nloyfer/wgbs_tools/blob/master/LICENSE.md. WGBS samples were downloaded from GSE186458.

## Funding

This work was supported by the Israel Science Foundation (grants no. 1250/18, 3099/22, 259/23). N.L is supported by the Center for Interdisciplinary Data Science Research (CIDR), and by the Leibniz Center for Research in Computer Science.

## Authors’ contributions

NL and TK conceived the idea. NL and JR implemented the software. TK supervised the work. All the authors contributed to the preparation of the manuscript. All the authors have read and approved the final manuscript.

## Supplemental tables

**Table S1.** The datasets used in Figure S1 to compare format sizes.

| Dataset | Method | Num of samples | Mean cover. | BAM size | bigwig size | pat size | Compression factor |
| --- | --- | --- | --- | --- | --- | --- | --- |
| Loyfer et al.[5] | WGBS | n=205 | 30x | 100.7Gb | 239.9Mb | 302Mb | **342.1** |
| Roadmap[30] | WGBS | n=91 | 32x | 127.6Gb | 248.7Mb | 256.4Mb | **488.1** |
| METABRIC[32] | RRBS | n=1,782 | 2x | 1.7Gb | 44.7Mb | 12.8Mb | **137.2** |
| Cheng et al.[33] | WGBS | n=52 | 1.5x | 5.1Gb | 136.4Mb | 45.5Mb | **113.7** |

**Table S2.**
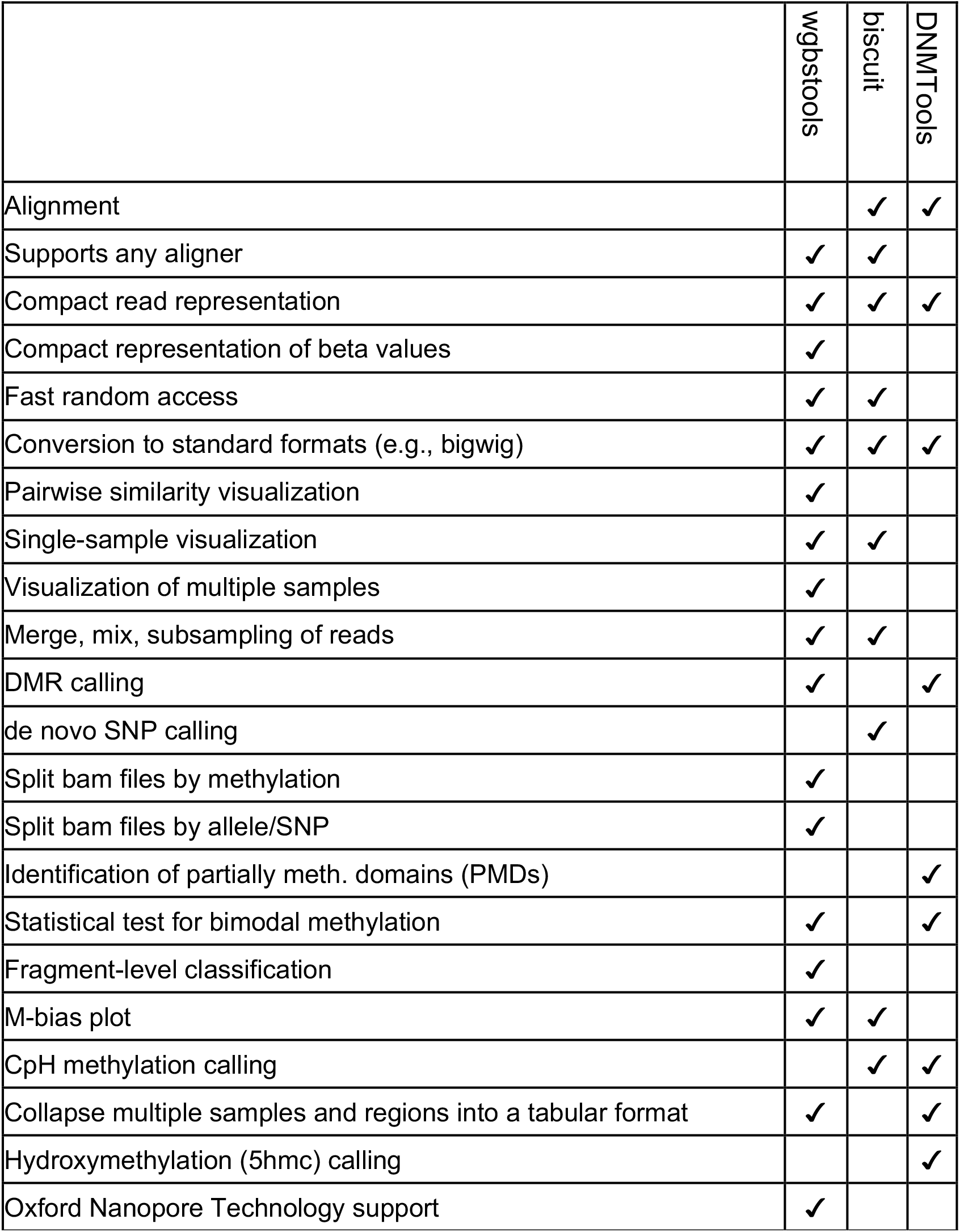
Comparison of available features in wgbstools and other software tools.

## Supplemental Figures

**Figure S1.**
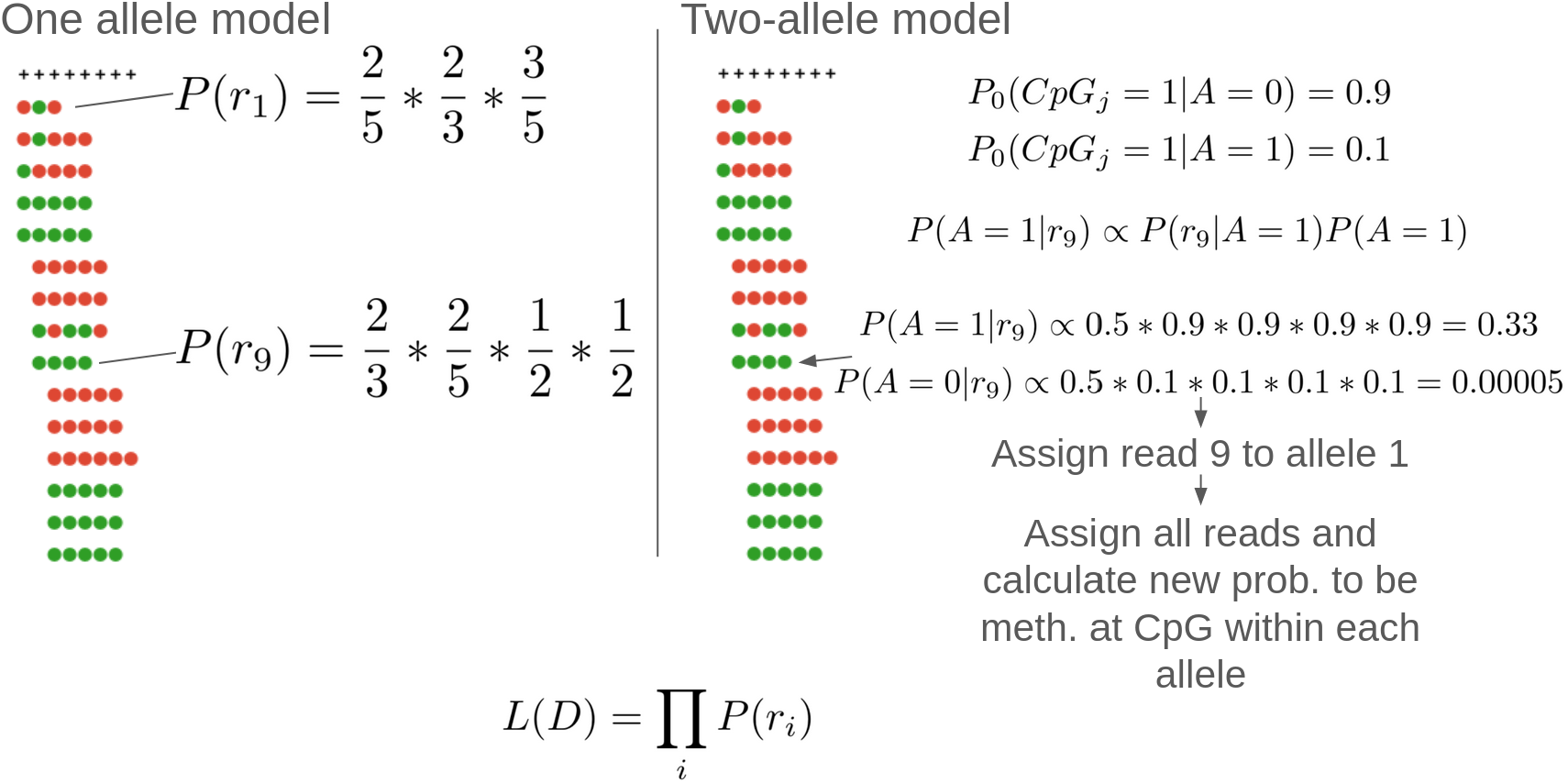
test_bimodal function. In the single allele model, for each CpG site we count the number of reads with a methylated CpG vs an unmethylated one. The first site shows two methylated and three unmethylated observances, yielding an estimated probability of ⅖ to be methylated. Likewise, the probability of the second site is ⅓, the third is ⅗, and the fourth and fifth are ½. Each read’s likelihood is thus the product of the probability to be methylated/unmethylated depending on the read’s data. In the two allele model we use an initial estimate of a probability to be methylated is 0.9 for allele 0 and 0.1 for allele 1. Thus, when calculating the probability of the allele being 1 for a fully unmethylated read is high, while for allele 0 it is very low. Each read’s allele assignment is determined using this calculation, and then the probability to be methylated on each allele for each CpG site is calculated, however, only using reads assigned to the allele. This is done iteratively, using expectation maximization, until the overall likelihood does not change.

## Notes

### Competing Interest Statement

The authors have declared no competing interest.

